# Microenvironment T-Type calcium channels regulate neuronal and glial processes to promote glioblastoma growth

**DOI:** 10.1101/2024.08.22.609229

**Authors:** Collin J. Dube, Ying Zhang, Shekhar Saha, Michelle Lai, Myron K. Gibert, Miguel Escalante, Kadie Hudson, Doris Wong, Pawel Marcinkiewicz, Ulas Yener, Yunan Sun, Esther Xu, Aditya Sorot, Elizabeth Mulcahy, Benjamin Kefas, Farina Hanif, Fadilla Guessous, Ashley Vernon, Manoj K. Patel, David Schiff, Hui Zong, Benjamin Purow, Eric Holland, Swapnil Sonkusare, Harald Sontheimer, Roger Abounader

## Abstract

**Background:** Glioblastoma (GBM) is the most common primary malignant brain tumor. The aim of this study was to elucidate the role of microenvironment and intrinsic T-type calcium channels (Cav3) in regulating tumor growth and progression.

**Methods:** We grafted syngeneic GBM cells into Cav3.2 knockout mice to assess the role of microenvironment T-Type calcium channels on GBM tumor growth. We performed single-cell RNA-seq (scRNA-seq) of tumors from WT and Cav3.2 KO mice to elucidate the regulation of tumors by the microenvironment. We used neurons from WT and Cav3.2 KO mice in co-culture with GBM stem cells (GSC) to assess the effects of Cav3.2 on neuron/GSC synaptic connections and tumor cell growth.

**Results:** Cav3.2 KO in the microenvironment led to significant reduction of GBM growth and prolongation of animal survival. scRNA-seq showed that microenvironment Cav3.2 regulates neuronal and glial biological processes. Microenvironment Cav3.2 downregulated numerous genes associated with regulating the OPC cell state in GBM tumors such as SOX10 and Olig2. Neuronal Cav3.2 promoted neuron/GSC synaptic connections and GSC growth. Treatment of GSCs with the Cav3 blocker mibefradil downregulated genes associated with neuronal processes. The Cav3 blocker drug mibefradil synergized with temozolomide (TMZ) and radiation to reduce in vivo tumor growth and prolong animal survival.

**Conclusions:** Together these data reveal a role for microenvironment Cav3 in promoting GBM tumor progression through regulating neuronal and glial processes particularly associated with the OPC-cell state. Targeting both intrinsic and microenvironment Cav3 with the inhibitor mibefradil significantly enhanced the anti-GBM effects of TMZ and radiation.

**Key Points:**

- Microenvironment Cav3.2 promotes GBM progression
- Microenvironment Cav3.2 promotes neuronal and glial processes
- Pharmacological targeting of intrinsic and microenvironment Cav3 synergizes with TMZ/radiation

**Importance of the Study:** In this study, we demonstrate for the first time that microenvironment Cav3.2 contributes to GBM progression and growth by regulating neuronal and glial processes. Our findings highlight the importance of T-type calcium channels in the microenvironment as well as the tumor and provides preclinically relevant data for the use of mibefradil to inhibit GBM growth in combination with standard of care therapies.

## Introduction

Glioblastoma (GBM), grade IV wildtype-IDH glioma, is the most common primary malignant brain tumor. GBM has a dismal prognosis of around 15 months despite aggressive standard of care therapy that includes surgery, chemotherapy and radiation [1, 2]. Gliomas have been shown to interact with the tumor microenvironment to promote growth and resistance to therapy. Gliomas form synaptic connections with neurons and neuronal activity drives growth and invasion through paracrine and electrical signaling [3-5] [6-9]. Gliomas also influence the neuronal circuits around them, increasing excitability and neuronal activity [10-15]. Neuronal signals from the tumor microenvironment influence neuronal gene expression and cell state of the tumor. For example, increased neuronal signaling in the tumor microenvironment promotes neuronal connections between oligodendrocyte precursor-like cells (OPCs) and neural precursor-like cells (NPCs) in the tumor with surrounding neurons [16, 17]. Importantly, GBM patients with high neural signatures exhibit decreased overall survival and progression free survival compared to patients with low neural signatures [18]. Gliomas utilize numerous mechanisms of neuronal interactions to collectively create a pro-tumor environment for growth, invasion and resistance to therapies. However, the exact nature and underlying mechanisms of neuron/GBM interactions and therapeutic strategies to disrupt these interactions have not been fully explored.

T-type calcium channels (Cav3) are a class low voltage gated calcium channels which are encoded by three genes (Cav3.1, Cav3.2, Cav3.3). Cav3 are expressed throughout the body, specifically within the central nervous system [19-23]. Cav3 are found both presynaptically and postynaptically throughout the central nervous system where they regulate presynaptic glutamate release as well as postsynaptic plasticity [23-25] [26]. Cav3.2 specifically has been shown to regulate neuronal firing in central nervous pathologies such as anxiety, memory disorders, epilepsy and chronic pain [27-30]. Our lab and others previously showed that Cav3 are overexpressed in GBM and regulate malignancy parameters such as growth, invasion, death and stemness [31-34]. However, the role of microenvironment Cav3 in GBM progression has not been investigated to date.

This study aimed to elucidate the role of microenvironment Cav3.2 on GBM tumor parameters including neuron/GBM interactions and malignancy, and to assess the therapeutic effects of targeting microenvironment and cell intrinsic Cav3. We hypothesized that loss of Cav3.2 within the brain tumor microenvironment would decrease tumor growth as well as neuron-GBM signaling processes. We grafted syngeneic GBM cells into whole body Cav3.2 KO mice to elucidate the effect on tumor growth, survival and proliferation. Loss of Cav3.2 in the microenvironment decreased neuronal and glial processes in the tumor, specifically those associated with the OPC cell state. Co-cultures of Cav3.2 KO neurons with GSCs showed decreased neuron/GBM projections and decreased tumor cell proliferation. Inhibition of Cav3 in GSCs significantly decreased genes associated with neuronal processes. Blockage of microenvironment and intrinsic Cav3 with the FDA-approved repurposed Cav3 blocker mibefradil sensitized tumors to temozolomide and radiation therapy. This is the first study elucidating the role of microenvironment Cav3.2 in promoting GBM tumor growth through regulating neuronal and glial processes.

## Methods

### Cell lines

The mouse GBM cell lines GL261 and CT2A were cultured in 500 mL Dulbecco’s modified eagle media (DMEM) (Gibco) containing 5 mL pen/strep, and 50 mL FBS. Glioblastoma Stem cell lines (GSCs) G34 were cultured in neurobasal (L-glutamine negative) media (Gibco, #.21103-049) containing 5 mL pen/strep, 5 mL B-27 without Vit-A (Gibco), 2.5 mL N-2 (Gibco), 1 mL EGF, 1 mL FGF, and 1.25 mL L-Glutamine. All cell media contained 5 μL Plasmocure reagent to prevent mycoplasma contamination.

### Orthotopic xenografting

For all xenografts male and female mice were used equally. For constitutive Cacna1h (Cav3.2) knockout studies, C57Bl6 wildtype (WT) and Cav3.2 knockout mice (The Jackson Laboratory) were used. Mice were anesthetized with ketamine and placed in a stereotactic apparatus. The cranium was exposed via a midline incision and 20,000 tumor cells (GL261 and CT2A) in 1 μl of sterile PBS was intracranially injected into the cortex; stereotactic coordinates used were as follows: 0.5mm lateral to midline, 1.0 mm anterior to bregma, -1.8 mm deep to the cranial surface. For drug studies in Nude athymic mice (Foxn1nu, Envigo), 100,000 tumor cells (G34) were injected in 1 μl of sterile PBS into the striatum; stereotactic coordinates used were as follow: 0.5 mm lateral to midline, 1.0 mm posterior to bregma, 2.5 mm deep from cranial surface. Brains were scanned with magnetic resonance imaging (MRI) on a 7 Tesla Bruker/Siemens ClinScan small animal MRI on the 18th day after cell transplantation. Tumor volumes were quantified using OsiriX Lite software (Version 12.5.2). For survival studies mice were monitored carefully and sacrificed when they displayed symptoms of tumor development including lethargy and head tilt.

### RCAS/TVA mouse model

For all RCAS/TVA mouse experiments, male and female mice were used equally. DF-1 cells were transfected with plasmids of RCAS-Scrambled, RCAS-shCav3.1, RCAS-Cav3.2, RCAS-PDGFB, and RCAS-Cre. Mice used in this experiment are the XFM model which is *Ntv-a Ink4a-Arf-/-LPTEN [35]*. Mice used for this experiment were 4 to 6.5 weeks old. Two microliters (1:1 mixtures of RCAS-PDGFB, RCAS-CRE and either RCAS-Scrambled, RCAS-shCav3.1, RCAS-shCav3.2) of 4 × 10^4^ transfected DF-1 cell suspensions were injected into the SVZ zone with the stereotactic coordinates 1.0 mm lateral to midline, 1.0 mm anterior to bregma, -2.5 mm deep from the cranial surface. Brains were scanned with a 7 Tesla Bruker/Siemens ClinScan small animal MRI on the 88th day after cell transplantation. Tumor volumes were quantified using OsiriX Lite software (Version 12.5.2). For survival studies mice were monitored carefully and sacrificed when they displayed symptoms of tumor development.

### Neuron-GBM co-cultures

Neurons were isolated from the brains of P1-P3 C57/BL6 WT, Cav3.2 or C57/BL6 Cav3.2 KO mice (The Jackson Laboratory) using the Neural Tissue Dissociation Kit-Postnatal Neurons (Miltenyi Biotec), followed by the Neuron Isolation Kit, Mouse (Miltenyi Biotec) as per the manufacturer’s instructions. After isolation, 300,000 neurons were plated on glass coverslips that were pretreated with poly-L-lysine for 1 hr at room temperature followed by 5 μg/ml mouse laminin for 3 hrs at 37C. Neurons were maintained in neurobasal medium (Invitrogen) supplemented with B27 supplement (2% v/v) and L-glutamine (0.5mM). After 5 days GSCs (50,000) were added and co-cultured in neurobasal medium. For EdU proliferation assays, co-cultures were plated for 48 hrs before treatment with EdU (10 μM) and incubated for a further 24 hrs. Following incubation, the cultures were fixed with 4% paraformaldehyde (PFA) for 20 mins at room temperature and stained using the Click-iT EdU cell Proliferation Kit (Thermo Fisher Scientific). The cells were then stained with mouse anti-nestin (1:500 Abcam), chicken anti-neurofilament (1:500, Aves Lab) overnight at 4°C. Following washing steps, the coverslips were incubated in secondary antibodies Alex 555 donkey anti-mouse IgG and Alex 647 donkey anti-chicken IgG and mounted using Prolong Gold Mounting medium (Life Technologies). Images were collected on a Leica Stellaris and the proliferation index was quantified at percentage of EdU-labeled GSCs (Nestin+) over total GSCs.

### Immunohistochemistry

Tumor-bearing mice were anesthetized with isoflurane inhalation followed by cervical dislocation followed by transcardial perfusion with 20 ml of PBS. Brains were dissected out and fixed in 4% PFA overnight at 4C. Afterwards brains were transferred to 30% sucrose. Brains were embedded in Tissue plus OCT (Thermo Fisher) then sectioned in the coronal plane at 10 μM using a cryostat (Thermo Cryostar NX50). For immunohistochemistry, coronal sections were incubated in a blocking solution (5% normal donkey serum, 0.3% Triton X-100 in PBS) at room temperature for 20 minutes. Chicken anti-GFP (1:500, abcam), rabbit anti-Ki67 (1:500, SantaCruz), rabbit anti-Neun (1:500, Abcam), mouse anti-Nestin (1:500, Abcam), mouse anti-Vglut1 (1:500, Abcam) were diluted in primary diluting solution (5% normal donkey serum, 0.3% Triton X-100 in PBS) and incubated overnight at 4°C. Sections were washed four times in PBT and incubated with Alexa 488 donkey anti-mouse, Alexa 555 donkey anti-rabbit, Alexa 647 donkey anti-chicken all at 1:250 (Thermo Fisher) in secondary diluting solution at room temperature for two hours. Sections were washed with PBT and mounted with ProLong Gold Antifade Mountant with DAPI (Thermo Fisher). Images were collected on a Leica Stellaris 8. Proliferation index was quantified at percentage of Ki67+ GFP+ cells over total GFP cells. Percentage of VGLUT1-GFP was calculated as VGLUT1+GFP+ over total GFP cells.

### Single cell RNA-Sequencing

Tumor-bearing mice were anesthetized under isoflurane inhalation followed by cervical dislocation. Brains were dissected under fluorescent microscope with GFP positive tumors isolated and placed on ice. One male and one female mouse was isolated from each group to control for sex-dependent variability. Tumors were mechanically and enzymatically dissociated using a papain-based brain tumor dissociation kit (Miltenyi Biotec). Samples were washed with PBS and red blood cell removal was performed as described (Miltenyi Biotec). Samples were next subjected to Debris removal as described (Miltenyi Biotec). Following this cells were stained with Live dead dye (7-AAD). Samples were subjected to FACS sorting (Influx Cell Sorter) for live cells. Samples were processed and sequenced using 10X Genomics according to validated standard operating procedures established by the University of Virginia Genome Analysis and Technology Core, RRID:SCR_018883.

### Single cell RNA-seq analysis

Sequencing files were downloaded for each flow cell lane and fastq files were merged. Reads were mapped to the mouse genome mm10 using 10X CellRanger (7.1.0). For single cell sequencing analysis standard procedures for filtering, mitochondrial gene removal, doublet removal, variable gene selection, dimension reductionality and clustering were performed using Seurat (version 5.0.2). A cell quality filter of greater than 200 features but fewer than 7500 features per cell and less than 25% of read counts attributed to mitochondrial genes was used. Dimensionality reduction was performed using principal component analysis and visualized with the Elbowplot function to find neighbors and clusters (resolution 0.3). Uniform manifold approximation and projection was performed for visualizing cells. Cell populations were identified using singleR with the celldex mouse RNA-seq data set as reference along with FindAllMarkers and the inferCNV (v1.18.1) algorithm to annotate clusters. Differentially expressed genes in tumors between WT and Cav3.2 KO were identified by FindMarkers using default settings. Gene Ontology Biological Pathways, Cellular Compartments and Molecular Functions of DEGS were performed using ClusterProfiler. Single cell RNA-Seq data can be found at the NIH GEO database (GSE273629).

Identification of co-expressed genes within the tumor cells was carried out using hdWGCNA (v0.2.23). Meta cells were constructed using harmony dimensionality reduction with 25 neighboring cells [36] using the function construct metacells. Modules were identified using the ConstructNetwork function (setDatExpr=FALSE) below the optimal soft threshold. To identify module feature genes, the ModuleEigenees function was used followed by computing the module connectivity to identify hub genes with the ModuleConnectivity function. To identify differentially expressed module eigengenes between WT and Cav3.2 KO, the FindDME function was used. Module scores representing the average expression of all the modules on the corresponding WGCNA modules were computed using the AddModuleScore function in Seurat. Pathway enrichment analysis of gene modules was performed using enrichR (v3.2).

Cell communication analysis was carried out using CellChat (v2.1.2). CellChat objects were created using createCellChat followed by inference of cell-cell communication using computeCommunProb. CellChat objects were created for both WT and KO samples then merged using MergeCellChat for comparison between the samples using the compare Interactions function. Differential number of interactions and interaction strength were visualized between WT and KO using NetVisual_heatmap. Dysfunctional signaling of Ligand-Receptor pairs was performed using IdentifyOverExpressedGenes then visualized based on sender and receiver cell types using NetVisual_bubble.

### Publicly available datasets

To examine expression of Cav3 in publicly available single cell datasets the Single Cell Portal was used [37]. Expression of Cav3 was queried in the Neftel dataset [17] and visualized by cell state using the single cell portal. Expression of Cav3 was queried in the Richards dataset [38] using the single cell portal. Expression of Cav3 was examined in publicly available spatial transcriptomic using the Ravi et al datasets [36] which is accessible using the SPATA2 tool box (v2.0.4).

### Statistical analysis

Comparisons between means of samples were performed using Students t-tests and one-way Anovas. All quantitative results are shown with means ± SEM. Molecular experiment tests, and computational experiment tests were performed using RStudio.

## Results

### Cav3.2 expression in the microenvironment promotes GBM progression and reduces survival

To elucidate the role of microenvironment Cav3.2 in glioma growth, we generated syngeneic GBM tumors using GL261 expressing Green fluorescent protein (GFP) into Cav3.2 KO (Cacna1h^-/-^) or WT C57/BL6 mice. After 21 days of engraftment of GBM cells, MRI scans to visualize and measure the tumors were performed. The MRI showed significant reduction in tumor volume in Cav3.2 KO mice compared to WT mice (Fig. 1A,B, Supp 1A,B). We next sought to determine if microenvironment Cav3.2 KO was sufficient to extend survival of syngeneic GBM tumor-bearing mice. Cav3.2 KO mice survived significantly longer than WT mice implanted in two syngeneic GBM models using GL261 and CT2A cells (Fig. 1C,D). Next, we assessed whether microenvironment Cav3.2 KO impacted tumor proliferation through histological analysis of Ki67 stained tumor sections. We found a significant reduction in the proliferation index in tumors in the Cav3.2 KO compared to WT mice (Fig 1E). Taken together, these findings indicate that microenvironment Cav3.2 promotes tumor growth and cell proliferation and decreases survival.

**Figure 1:**
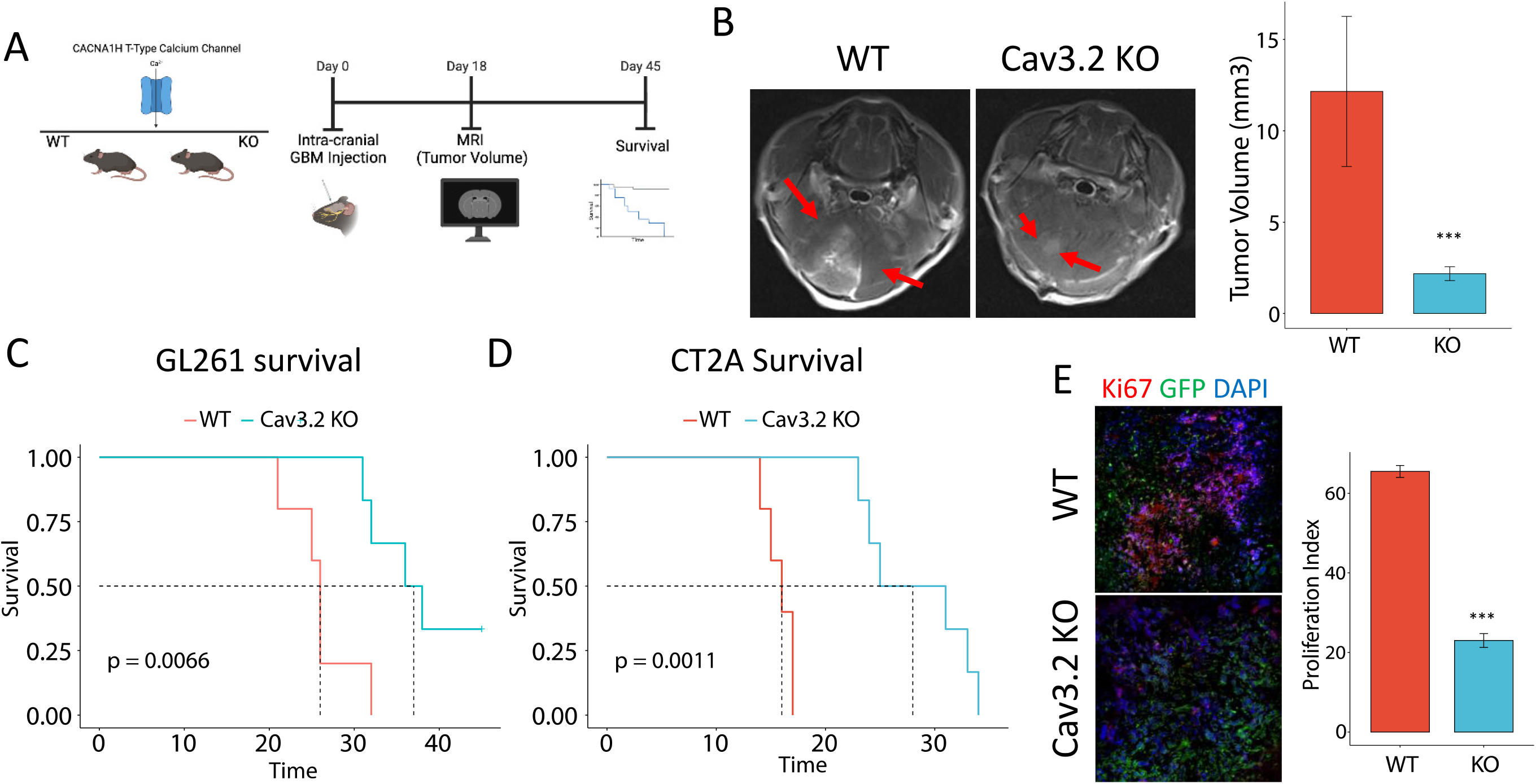
Microenvironment Cav3.2 promotes tumor progression and reduces survival. A) Schematic of experimental workflow for WT and Cav3.2 KO experiments. B) Representative MRIs and quantification showing that microenvironment Cav3.2 KO leads to significant reduction in GBM tumor volume. Red arrows indicate the tumor location. C & D) Kaplan Meir survival curves for WT and Cav3.2 KO mice in GL261 and CT2A xenograft tumors respectively. E) Representative immunofluorescence staining of Ki67 in WT and Cav3.2 KO with quantification showing a significant reduction in proliferation index. ***<0.001

### Microenvironment Cav3.2 regulates genes important for neuron-GBM interactions

To molecularly define the effects of microenvironment Cav3.2 on GBM tumors we performed single-cell RNA-seq (scRNA-seq) on syngeneic tumors implanted in WT and Cav3.2 KO mice (Fig. 2A). We utilized established markers as well as the infercnv package to differentiate microenvironment populations from tumor cells (Fig. 2B, Supp 2A-H). We performed differential gene expression analysis of the WT and Cav3.2 KO tumors to identify the downregulated genes in the tumor and assess how loss of Cav3.2 in the microenvironment impacts tumor signaling. To gain insight into what the downregulated genes were doing we performed Gene Ontology (GO) Biological pathway analysis which showed downregulation of nervous system processes including gliogenesis, glial cell differentiation, regulation of synapse organization, structure or activity and axon guidance in GBM cells in the Cav3.2 KO compared to WT tumors (Fig. 2C,D, Supp 3A,B). Gene ontology cellular compartment analysis of the downregulated genes showed enrichment of genes associated with neuronal processes such as postsynaptic specialization, neuron to neuron synapse, asynaptic synapse, postsynaptic density and distal axon (Fig. 2 E,F). Analysis of the upregulated genes showed enrichment for biological pathways associated with positive immune response, phagocytosis and T-cell activation (Supp 3C,D). These data indicate a role of microenvironment Cav3.2 in regulating neuronal (synapse organization, axon guidance) and glial processes (gliogenesis, glial cell differentiation) in the tumor, which are important for regulating connections of the tumor to the other cell types such as neurons and astrocytes.

**Figure 2:**
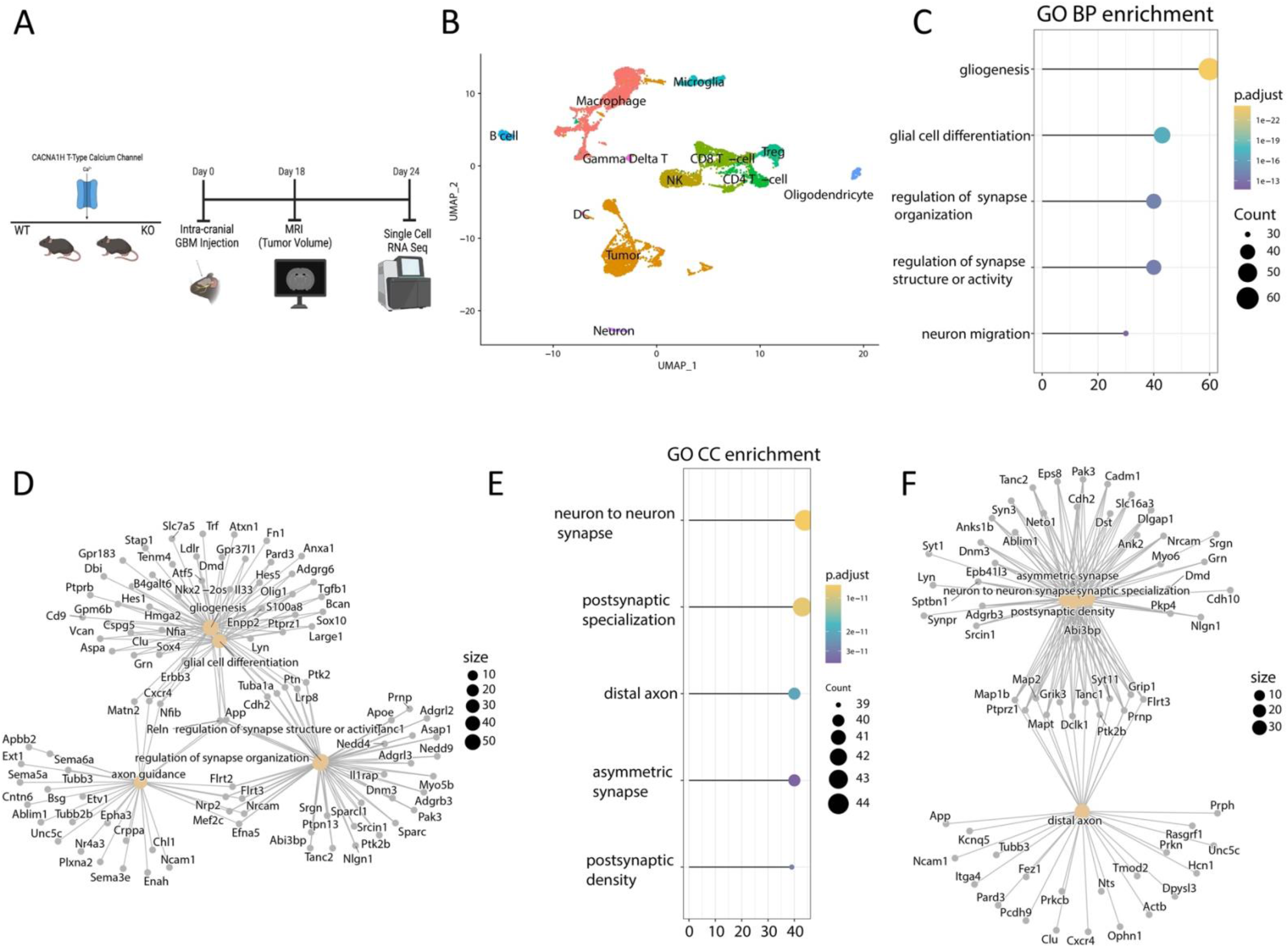
Microenvironmental Cav3.2 regulates neuronal and glial processes. A) Schematic showing the experimental workflow for single-cell RNA-seq tumor isolation. B) UMAP of the distinct cell types isolated from WT and KO mice. C) Gene ontology Biological Pathways of downregulated genes in Cav3.2 KO tumors. D) Cnetplot of GO BP pathways and associated genes. E) Gene ontology Cellular Compartment of downregulated genes in KO tumors. F) Cnetplot of GO CC pathways and corresponding genes.

### Microenvironmental Cav3.2 regulates OPC-related genes and cell states in GBM cells

To gain further insight into the role of microenvironment Cav3.2 in the context of GBM, we utilized single cell weighted gene co-expression network association analysis (hdWGCNA) on tumor cells from WT and Cav3.2 KO mice (Supp 4A,B). This analysis identifies modules of co-expressed gene regulatory networks across multiple cell clusters. Gene ontology of the tumor modules identified numerous biological pathways associated with the specific modules such as antigen receptor mediated signaling pathways and synaptic pruning (Supp 4C). After identifying the co-expression modules, we performed differential module eigengene (DME) analysis to examine which modules are upregulated and downregulated in Cav3.2 KO tumors (Supp 4D). The analysis identified tumor module 4 as downregulated in tumors from Cav3.2 KO mice. The most co-expressed genes in this module form an interconnected graph and consist of genes associated with oligodendrocyte differentiation (PTPRZ1, SOX6, SOX10) and nervous system development (RBFOX2, NAV3, S100B, GPM6B, ADGRL3) based on GO (gene otology) pathway analysis (Fig 3A,B). Additionally analysis of the tumor 4 module and differentially expressed genes revealed an overlap of genes associated with the OPC cell state (Supp 4E,F). We examined the OPC cell state and found a decreased enrichment of the OPC cell state in Cav3.2 KO compared to WT microenvironment tumors (Fig 3C). We then characterized the tumors by cell state according to neftel, which demonstrated that tumors from Cav3.2 KO microenvironment exhibit a decrease in OPC cell states and a shift towards an increase in AC cell state (Supp 5A). We next assessed the differential regulon activity of tumors, which revealed a decreased activity of Olig2 and Sox10 in OPC and NPC cell states (Supp 5B). The decreased enrichment of the OPC cell state as well as activity of OPC related transcription factors led to the examination of SOX10 and Olig2 specifically, which showed downregulation in tumors in the Cav3.2 KO microenvironment (Fig 3D,E, Supp 5C). The OPC cell state has been shown to interact with neurons in the tumor microenvironment [16]. Therefore, we examined publicly available spatial transcriptomic data by Ravi et al [36] and found that enrichment of SOX10 and Olig2 in the transition and infiltration region of the tumors specifically OPC-like cells indicative of connections with neurons in the microenvironment (Fig 3F, Supp 5D). The cell state of GBM regulates the ability of tumor cells to interact with the various cell types of the tumor microenvironment. Our data suggest that microenvironment Cav3.2 regulates the OPC cell state of the tumor through SOX10 and Olig2 which are important transcription factors for regulating genes responsible for the OPC-like cell state.

**Figure 3:**
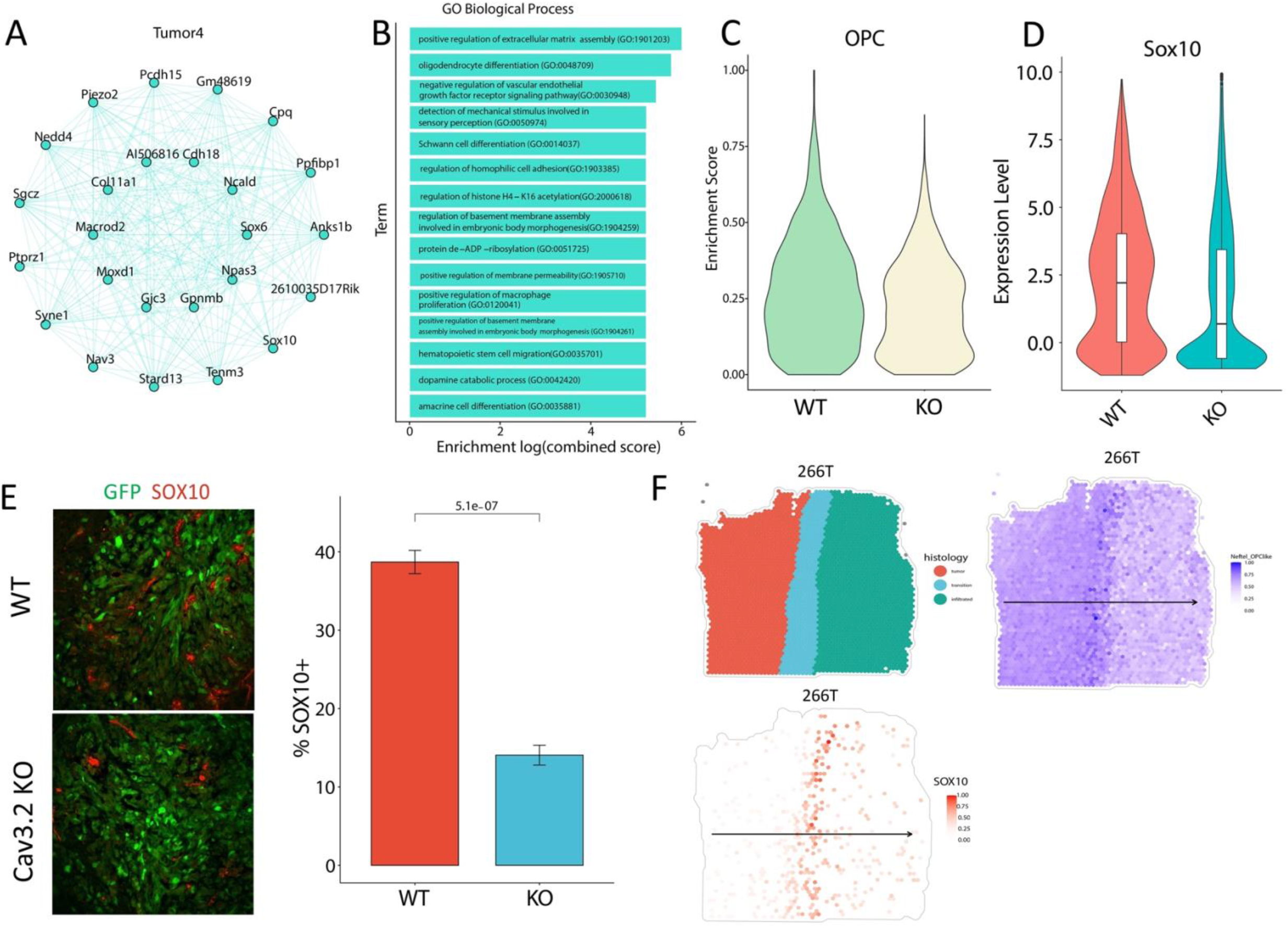
Cav3.2 KO downregulates OPC cell state genes. A) Network graph of the top 25 co-expressed genes in the Tumor 4 module of the hdWGCNA analysis. B) Gene Ontology Biological Pathways for the Tumor module 4. C) Enrichment score of the OPC cell state in WT and Cav3.2 KO tumors. D) Expression of Sox10 in WT and Cav3.2 KO tumors showing decreased expression. E) Representative immunofluorescent staining of SOX10 (Red) in WT and Cav3.2 KO tumors (GFP) with quantification of %SOX10 positive cells on the right. F) Spatial transcriptomic data of 266T showing the different zones of tumor infiltration (Left), OPC-like cells (Right) and Sox10 expression (Bottom) demonstrating enrichment of Sox10 in the OPC-like cells in the transition zone.

### Neuronal Cav3.2 enhances GBM growth and guides neuron/GBM connections

The downregulation of neuronal genes and neuronal processes in tumors from Cav3.2 KO led us to examine the role of Cav3.2 in neurons in the GBM microenvironment as Cav3.2 is expressed in glutamatergic neurons. To elucidate the role of Cav3.2 in neurons, we isolated neurons from WT and Cav3.2 KO mice for co-culture experiments with GBM stem cells (GSCs). The co-culture of WT neurons with GSCs enhanced the proliferation index of the GSCs (Fig. 4A). The enhanced proliferation was abolished when the GSCs were co-cultured with neurons isolated from Cav3.2 KO mice (Fig. 4A). To demonstrate that the proliferation effect was coming specifically from the neurons we utilized tetrodotoxin (TTX) to inhibit the neuronal activity which abrogated the additive effect of WT neurons on GSC growth (Fig. 4A). Next, we assessed whether direct contact was necessary to attenuate the growth effect. To this end, we applied conditioned medium from WT and Cav3.2 KO neurons to GSCs and found that this mirrored the results of the co-culture experiments (Fig. 4B). We also assessed the connections between WT and Cav3.2 KO neurons with GSCs in co-culture and found that Cav3.2 KO neurons exhibited less projections toward GSC cells compared to WT neurons (Fig. 4C). We assessed neuron innervation in WT and Cav3.2 KO tumors which showed significant decrease in glutamatergic neurons (Vglut1) present in the Cav3.2 KO tumors compared to WT tumors (Fig. 4D). This observation is consistent with the single cell Gene Ontology analysis which showed decreases in genes associated with axon guidance. To gain insight into how neurons affect GBM cells, we utilized CellChat to examine the ligand-receptor relationships between neurons and tumor cells. Examination of neurons-tumor showed decreased signaling associated with the glutamate pathway while upregulating GABA signaling in the Cav3.2 KO microenvironment (Supp 5E, 6A-C). These data demonstrate the importance of neuronal Cav3.2 in regulating GSC growth as well as neuron/GBM connections.

**Figure 4:**
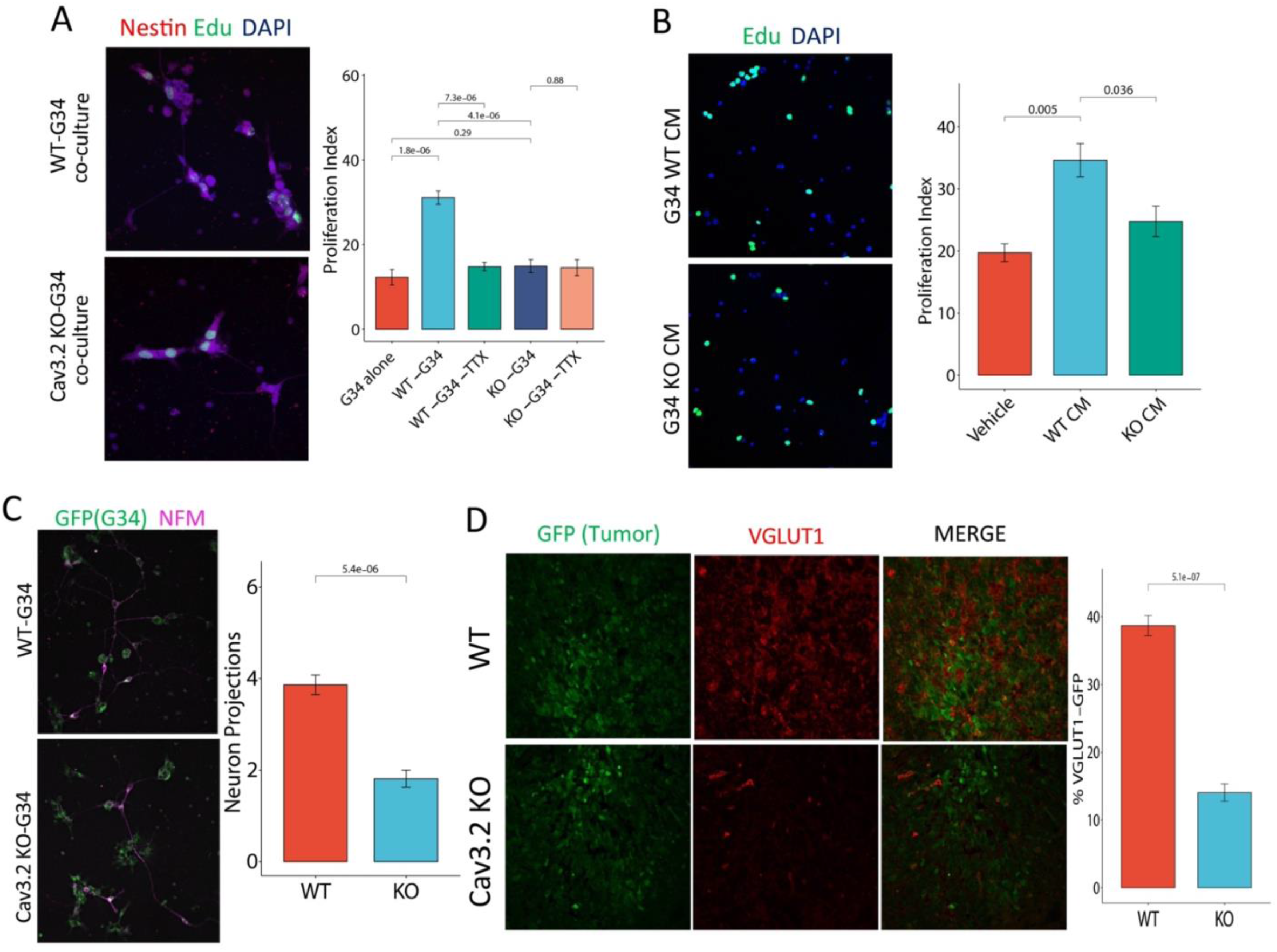
Neuronal Cav3.2 promotes GSCs growth and neuronal connections. A) Representative immunofluorescent images of Glioma Stem cells G34 (Red) co-cultured with WT and Cav3.2 KO neurons. Proliferative cells are labeled with EDU (Green) with quantification of proliferation index to the right. B) Representative immunofluorescent images of G34 cells subjected to WT and Cav3.2 KO Conditioned medium with quantification of proliferation index to the right. C) Representative immunofluorescent images of Glioma Stem cells G34-GFP (Green) co-cultured with WT and Cav3.2 KO neurons labeled with Neurofilament (Pink) with quantification of neuron projections to the right. D) Representative immunofluorescent staining of VGLUT1 (Red) in WT and Cav3.2 KO tumors (GFP) with quantification of percent VGLUT1-GFP positive cells on the right.

### Therapeutic targeting of both tumor intrinsic and microenvironment Cav3 inhibits GBM growth

We next assessed the therapeutic value of targeting tumor cell intrinsic and microenvironment Cav3 in GBM. We previously showed a role for tumor intrinsic Cav3.2 in promoting GBM growth using a xenograft model [31]. To validate and expand these findings, we used the RCAS/TVA mouse model to deliver shRNA for Cav3.1 and Cav3.2 to GBM tumors generated by implanting vector infected chicken fibroblast cells (DF-1) into the SVZ of the XFM mouse model(*Ntv-a Ink4a-Arf-/-LPTEN*)[35]. Mice received DF-1 cells with RCAS-PDGFB, RCAS-Cre and either RCAS-Scrambled or RCAS-shCav3.1, RCAS-shCav3.2. We found significant reduction in tumor volume as well as significantly prolonged survival of mice in which tumor Cav3 was knocked down as compared to control mice (Supp 7A-D). We then assessed the effects of targeting both tumor intrinsic and microenvironment Cav3 on tumor growth using the repurposed FDA approved Cav3 blocker mibefradil that blocks all three Cav3 isoforms in humans and mice [39]. We tested the effects of mibefradil in combination with GBM standard of care therapy consisting of temozolomide (TMZ) and radiation. GSCs were stereotactically implanted into the striata of immunodeficient mice [2]. Six days after implantation mibefradil (24 mg/kg body weight) was administered by oral gavage every 6 hours for 5 days. TMZ (100 mg/kg body weight) was concurrently administrated via IP once a day for 4 days. Radiation (70 cGy) was administered in a small animal irradiator (Rad Source RS-2000) once a day for 4 days. MRI scans were performed 18 days after surgery and tumor volume was assessed. The effect of mibefradil alone significantly reduced tumor volume compared to control (Fig. 5 A,B). The triple combination of mibefradil with temozolomide and radiation significantly reduced tumor volume compared to the TMZ/radiation only group suggesting mibefradil gives an additive effect (Fig. 5 A,B). These data suggest that the inhibition of Cav3 in the microenvironment and in tumors enhances standard of care GBM therapy.

**Figure 5:**
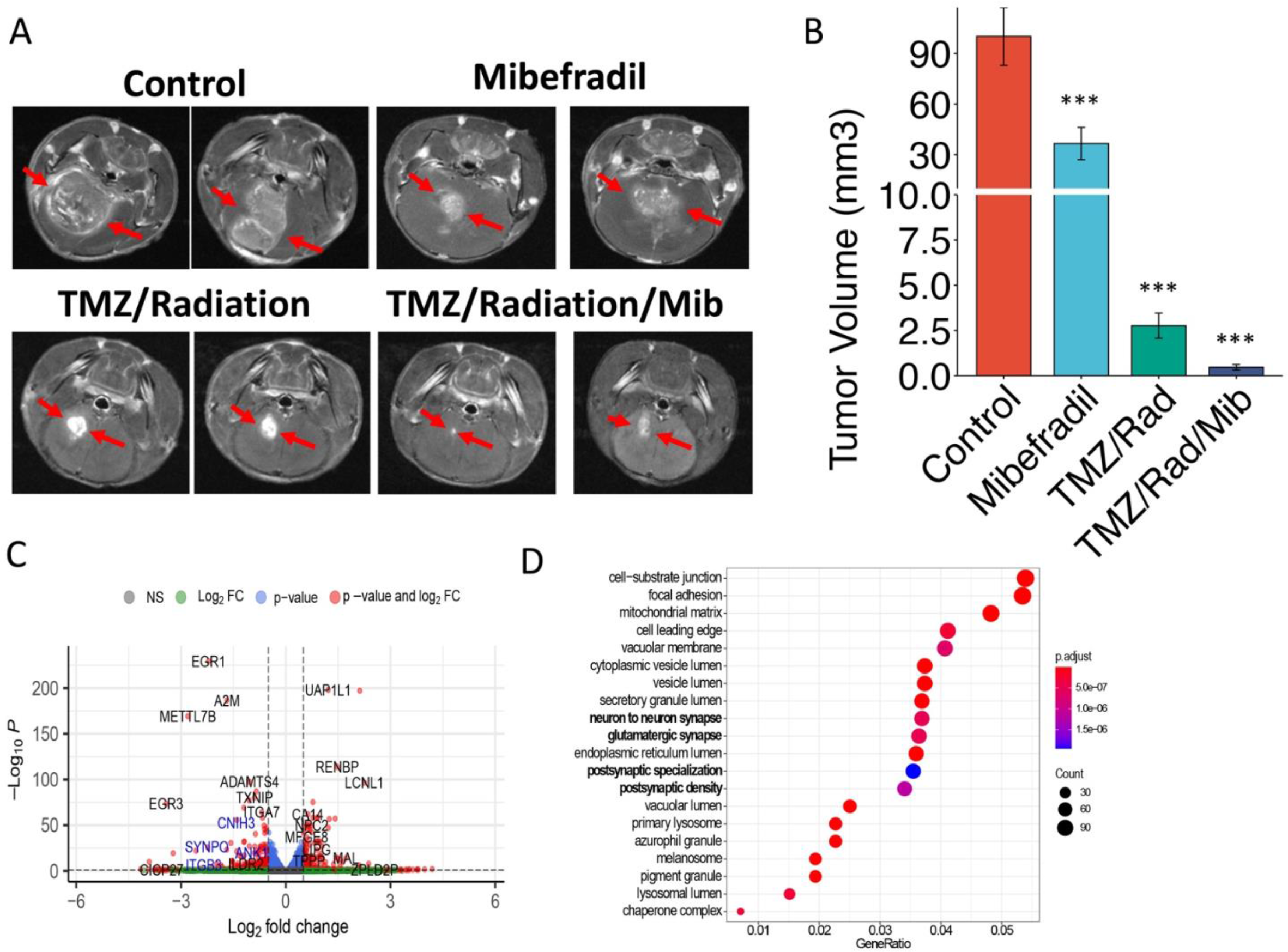
Mibefradil synergizes with standard of care to reduce tumor volume and downregulate neuronal processes. A) Representative MRIs of Vehicle, Mibefradil, TMZ + Radiation, Mibefradil + TMZ+ Radiation. B) Quantification of tumor volume. C) Volcano plot of the differentially expressed genes from GSCs treated with mibefradil. D) Dotplot of the Gene ontology Biological Pathway for the deregulated genes upon GSC treatment. ***<0.001

### The therapeutic effects of targeting Cav3 are associated with inhibition of neuronal processes

To confirm that the therapeutic effects of Cav3 inhibition are associated with the alteration of neuronal processes, we treated GSCs with mibefradil and performed RNA-sequencing. Mibefradil treatment downregulated key neuronal genes such as Synpo, Syn1, Vgf, Sema7a, Nxph4 (Fig. 5C). Analysis of differentially expressed genes revealed enrichment of biological processes associated with neuronal processes such as glutamatergic synapse, postsynaptic density and neuron to neuron synapse (Fig. 5D). The mibefradil data gave a similar effect as the Cav3.2 KO signle-cell RNA-seq data in downregulating genes associated with neuronal processes. We also analyzed TCGA data and found that Cav3 are co-expressed with genes associated neuronal processes such as axon Sema6a, Sema6c, Plxna2, Kcnc1, Grin2b, Shank2, Shank3, which are associated with neuronal processes such as axon guidance, synapse organization, neuron project and ionotropic glutamate receptor binding (Supp 8A,B). We also analyzed publicly available single cell RNA-seq [17] and found that Cav3.2 is enriched in NPC-like GBM cells (Supp 8C). We also found that Cav3.2 are enriched in the developmental transcriptional signatures compared to injury response signatures within the Rchards dataset (Supp 8D), which is important as the developmental signatures are more likely to interact with the neurons [38]. Additionally, analysis of publicly available spatial transcriptomic data of GBM shows enrichments of Cav3 in the NPC cell state and neuronal development spatial transcriptional program (Supp 8e) [36]. The above data demonstrate that Cav3 are co-expressed with neuronal genes and that the inhibition of Cav3 with mibefradil downregulates genes important for neuron-GBM processes.

## Discussion

Cell state is important for receiving signals from the microenvironment with the NPC/OPC cell states previously shown to form synaptic connections with tumors promoting growth and infiltration [16]. GBM tumor microtubules, which have been shown to connect to neurons, are more resistant to chemotherapy and radiation [40, 41]. Patients with high neural signatures exhibit decreased overall survival and progression free survival in GBM and high-grade gliomas [18]. The mechanism of neuron-tumor interaction is still understudied and requires further elucidation of both the neurons and tumors. Cav3 are expressed in glutamatergic neurons and regulate glutamate release as well as glutamate receptor plasticity [25]. Our study aimed to elucidate the role of Cav3 in the microenvironment and its effects on tumor growth.

We investigated the role of Cav3.2 in the GBM microenvironment by first utilizing whole body knockout mice which revealed that loss of Cav3.2 the microenvironment inhibited tumor growth, proliferation and increased survival. To gain insight into the mechanism of inhibition of tumor growth we performed single cell RNA-sequencing, which revealed that loss of Cav3.2 in the microenvironment decreased expression of genes associated with neuronal and glial processes such as gliogenesis, axon guidance and regulation of synapse organization. Among the downregulated genes were genes associated with regulating the OPC cell state such as Sox10 and Olig2 which had decreased regulon activity in Cav3.2 KO tumors. Sox10 is a transcription factor that is involved in development shifting neural crest cells toward oligodendrocyte precursor cells [42, 43]. In glioma Sox10 has been shown to induce gliomagenesis, regulate oligodendrocyte lineage and suppress astrocyte lineage [44, 45]. Sox10 has also been identified as a master regulator of the RTK1 epigenetic subtype which is associated with proneural signatures and loss of Sox10 shifts GBM cells towards a mesenchymal like state [46]. Microenvironmental Cav3.2 regulating OPC cell state as well as neuronal and glial processes points toward its role in regulating tumor connections to neurons. To validate these findings, we examined the role of neuronal Cav3.2 using neuron/GBM co-cultures and found that loss of Cav3.2 in the neurons led to a significant decrease of GSC proliferation in co-cultures as well as in conditioned media from neurons. Additionally, Cav3.2 KO neurons exhibited less neuronal projections towards GSCs. Additionally, Cav3.2 KO tumors showed less glutamatergic neuron innervation compared to WT tumors. Analysis of the single cell RNA-seq data with CellChat, which is used to infer cell-cell communication, showed that in the Cav3.2 KO microenvironment neuron-tumor communication is associated with a decrease in glutamatergic signaling and an increase in GABAergic signaling, which can inhibit glioma growth [47].

By analyzing publicly available single cell and spatial transcriptomic datasets we also found that Cav3 are enriched in NPC-like and OPC-like cell states as well as neurodevelopmental transcriptional signatures. Additionally, Cav3 was positively correlated with genes associated with neuronal processes. The inhibition of Cav3 downregulated neuronal associated genes specifically for processes associated with synapse formation and communication. The combination of mibefradil, temozolomide and radiation significantly decreased tumor volume compared to TMZ/radiation alone, presumably via the inhibition of both tumor intrinsic and microenvironment Cav3. These data support the use of mibefradil to complement standard GBM therapies and warrant testing this combination in clinical trials. The Cav3 inhibitor mibefradil has been examined in clinical trials in recurrent gliomas demonstrating safety and potential activity in combination with temozolomide [48]. Mibefradil was also assessed in a clinical trial in combination with radiation therapy in recurrent GBM patients which demonstrated safety and potential activity with radiation therapy [49]. Recent studies have modified mibefradil to improve the drug to create better radiosensitizers for treatment of GBM [34]. Our data along with the clinical trial data suggest that a phase 2 trials in newly diagnosed GBM patients is warranted.

In conclusion, our results represent the first characterization of Cav3.2 in the GBM microenvironment. Microenvironment Cav3.2 promotes GBM growth, proliferation and decreased survival. Microenvironmental Cav3.2. regulates GBM neuronal and glial cell processes and regulates OPC cell state within the tumor through Sox10. Neuronal Cav3.2 promotes glioma stem cell proliferation and neuronal connections. Within GBM tumors Cav3 channels are enriched in neurodevelopmental transcriptional states and correlated with neuronal genes. Blockage of Cav3 enhances the effects of standard GBM therapies most likely by altering tumor intrinsic as well as microenvironmental factors, supporting evaluation of mibefradil in future clinical trials.

## Authorship

Designing research studies: CD and RA. Conducting experiments and acquiring data: CD, YZ, SS, ML, MG, ME, KH, DW, PM, UY, YS, EX, AS, BK, FH, FG, AV. Analyzing Data: CD, RA.

Conceptual input: CD, RA, MKP, DS, HZ, BP, SS HS. Writing manuscript: CD and RA. Reviewing and editing of manuscript: All authors

