## Supplemental Fig for "Microenvironment T-Type calcium channels regulate neuronal and glial processes to promote glioblastoma growth"

### Supplemental Figures

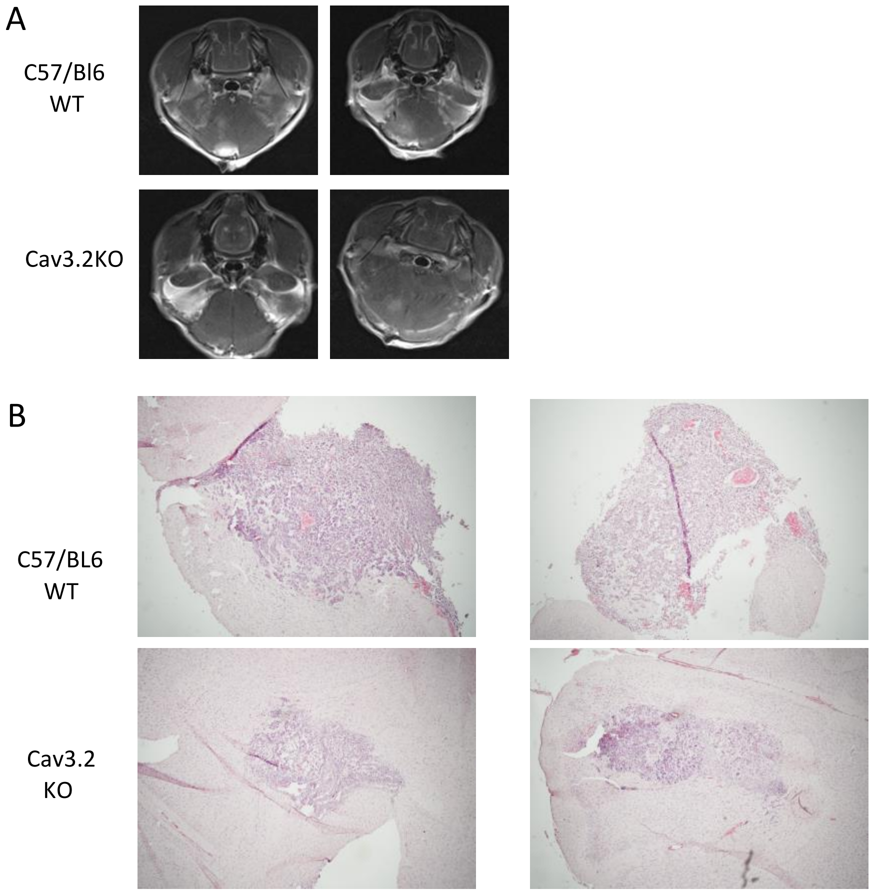


**Supplemental Figure 1**: A) Representative MRI images of GL261 xenografts in WT and Cav3.2KO mice. B) Representative H&E staining of tumor sections from GL261 xenografts from WT and Cav3.2 KO mice.

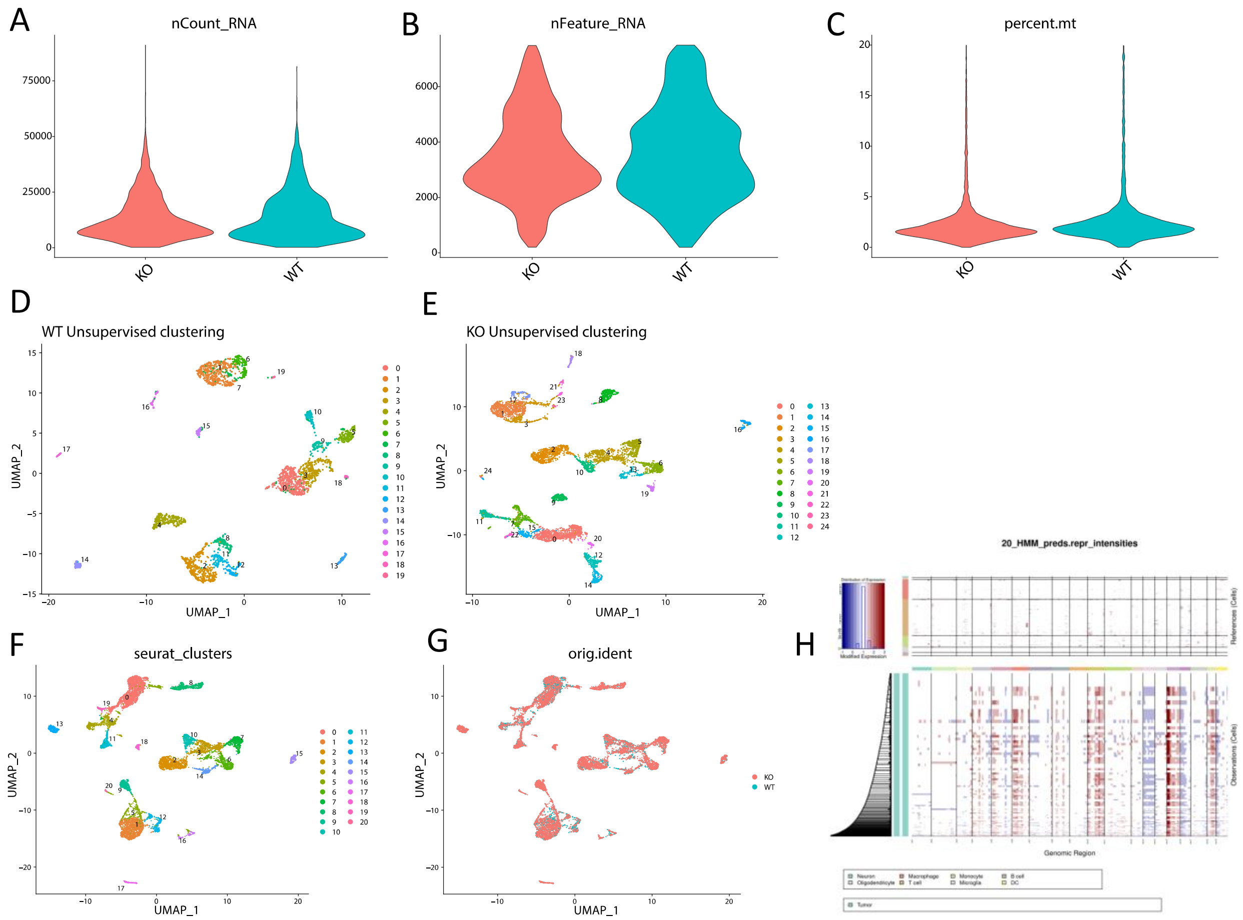


**Supplemental Figure 2:** Single-cell RNA sequencing of Wild type (WT) and Cacna1H-/- (KO) tumors. A-C. Quality control of sequenced tumors from WT and KO tumors. D,E) Unsuperivsed clustering of WT and KO tumors. F Seurat clusters for the merged object of WT and Cav3.2 KO tumors. G) UMAP showing the overlap of WT and Cav3.2 KO tumors in the merged object. H) Heatmap of inference of chromosomal CNAs with the top rows being non-malignant cells followed by tumor cells.

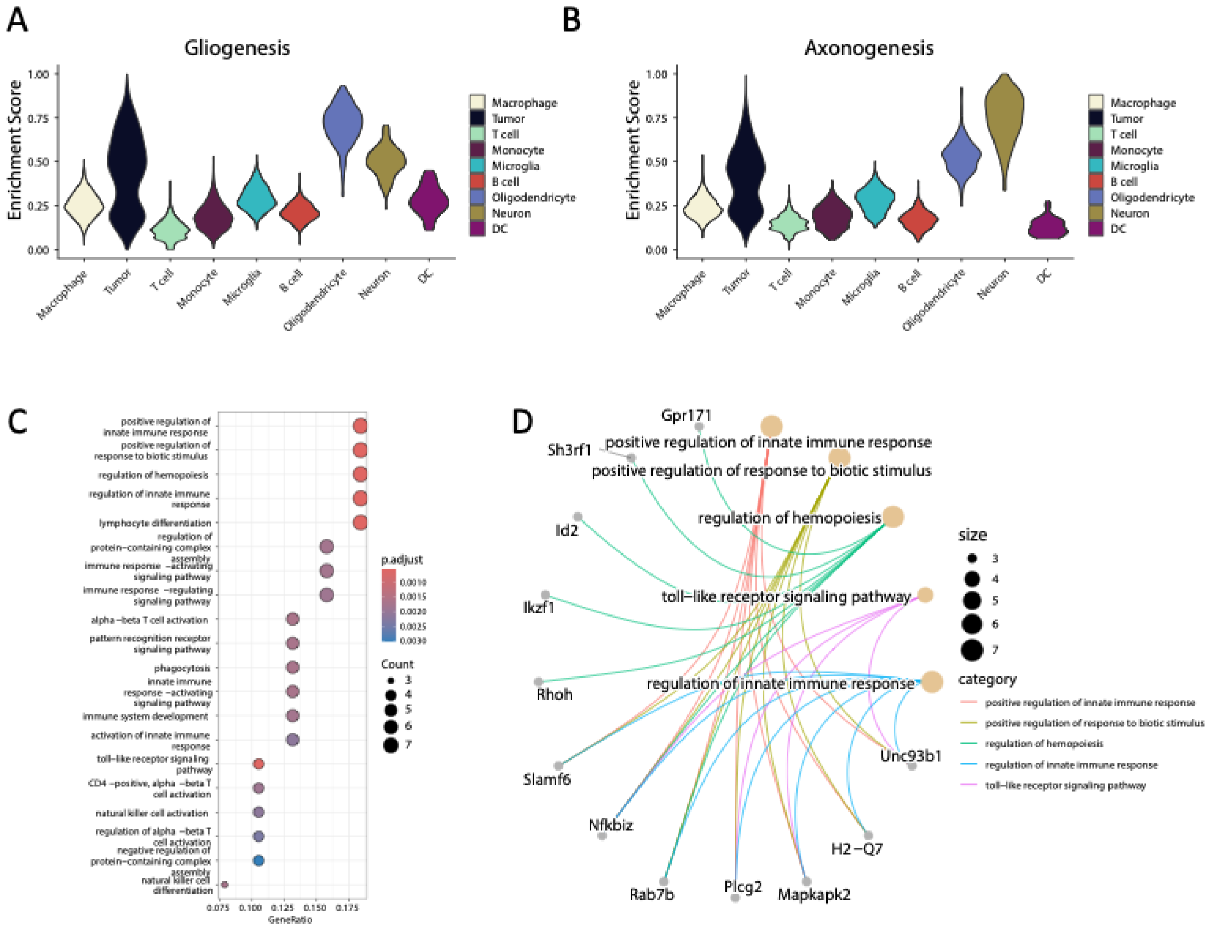


**Supplemental Figure 3:** A) Enrichment score of Gliogenesis genes across cell types. B) Enrichment score of Axongenesis genes across cell types. C) Gene Ontology Biological pathway analysis of upregulated genes in Cav3.2 KO tumors. D) Cnetplot of upregulated biological pathways and corresponding genes with each process.

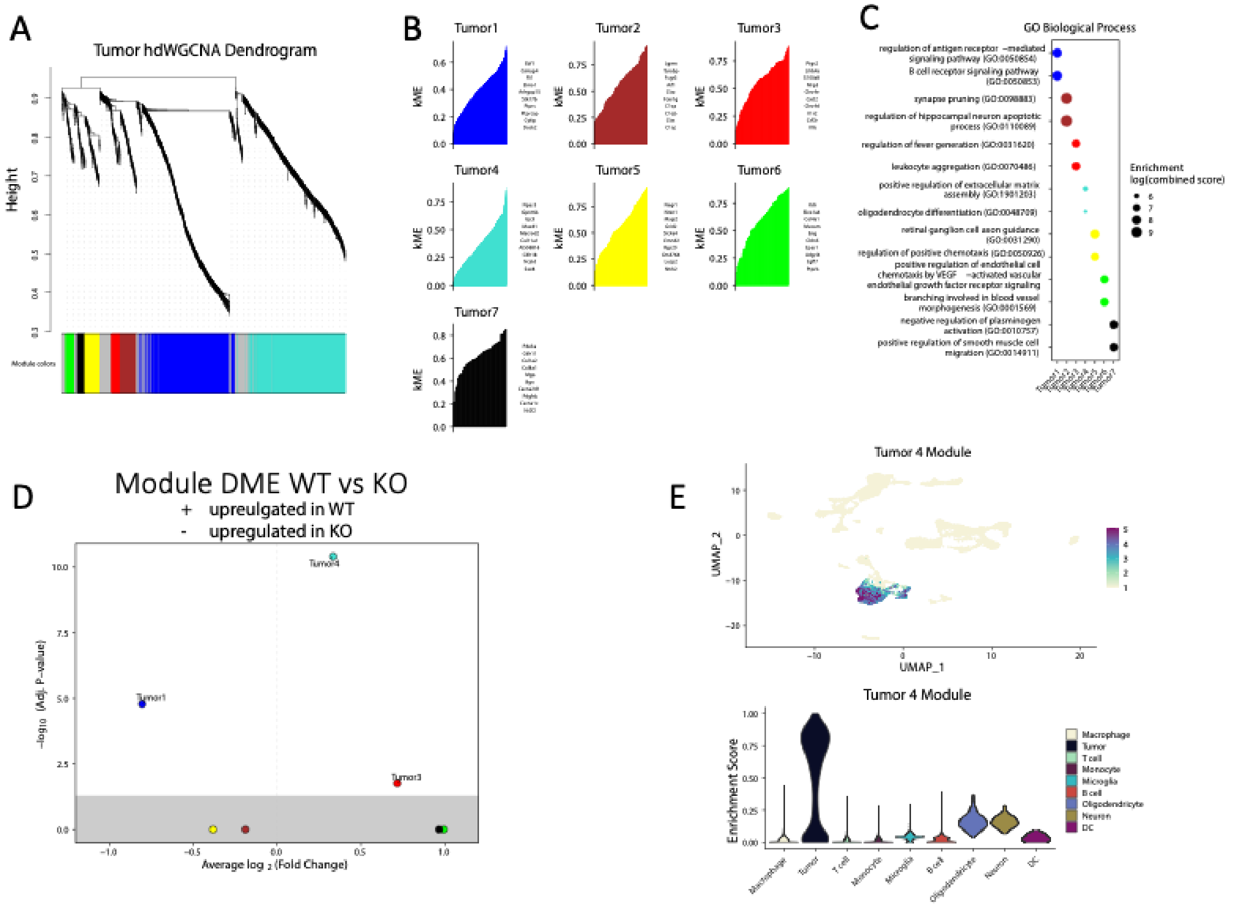


**Supplemental Figure 4:** A) Dendogram of co-expression modules single cell RNA seq tumors. B) kME for the tumor modules identified in hdWGCNA. C) Gene ontology biological processes for each of the tumor modules. D) Volcano plot of the differential module eigene between WT and Cav3.2 KO tumors. E) Feature plot showing overlay of tumor 4 module with corresponding Violen plot showing enrichmnet of the tumor 4 module in the cell types.

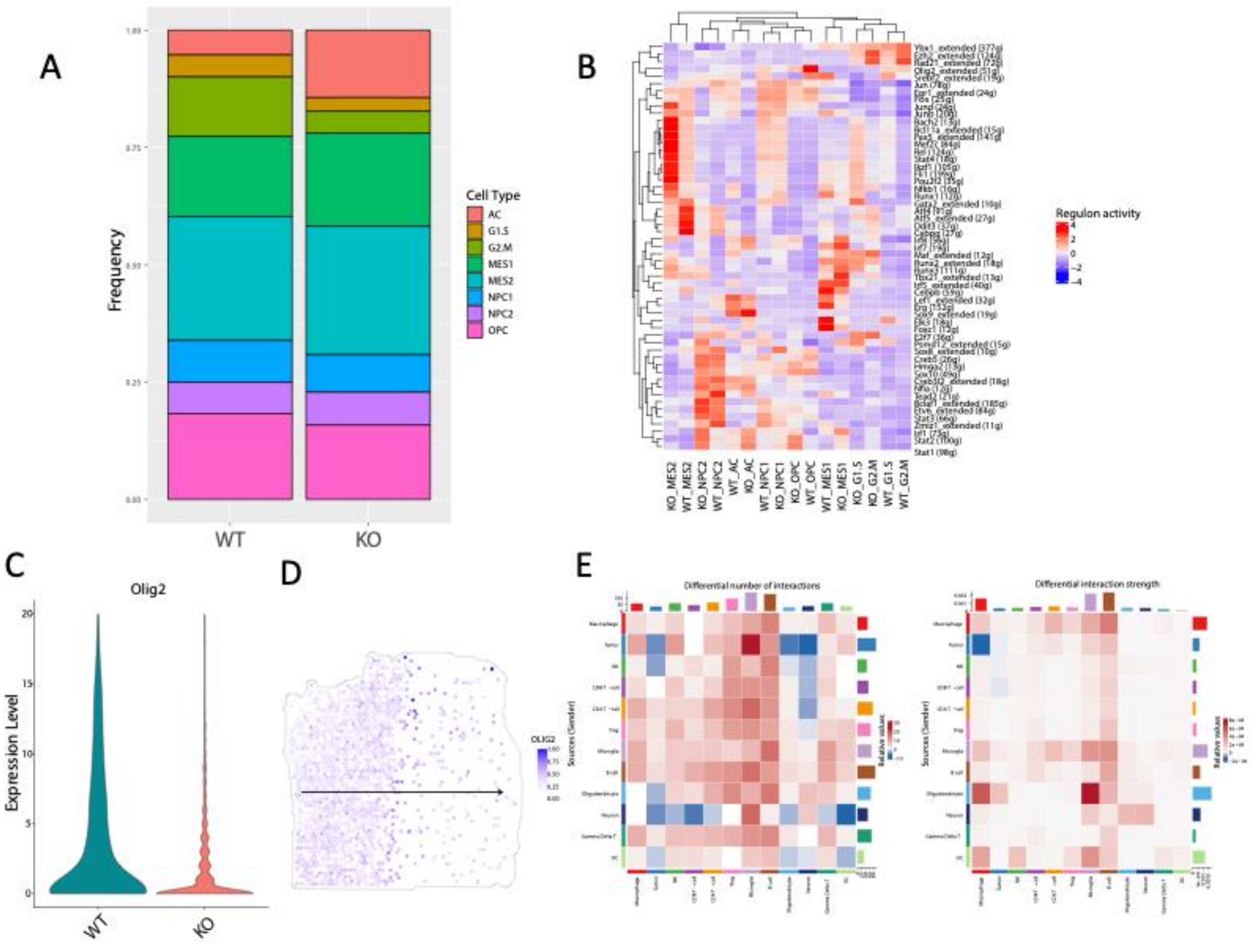


**Supplemental Figure 5:** A) Frequency of cell states in WT and Cav3.2 KO Tumors. B) Heatmap of regulon activity of transcription factors from Scenic analysis for each cell state within WT and Cav3.2 KO tumors. C) Expression of Olig2 in WT and Cav3.2 KO tumors showing decreased expression. D) Spatial transcriptomic data of 266T showing the Olig2 expression. E) Heatmap of differential number of interactions and interaction strength of cell senders and receivers between WT and Cav3.2 KO cellchats.

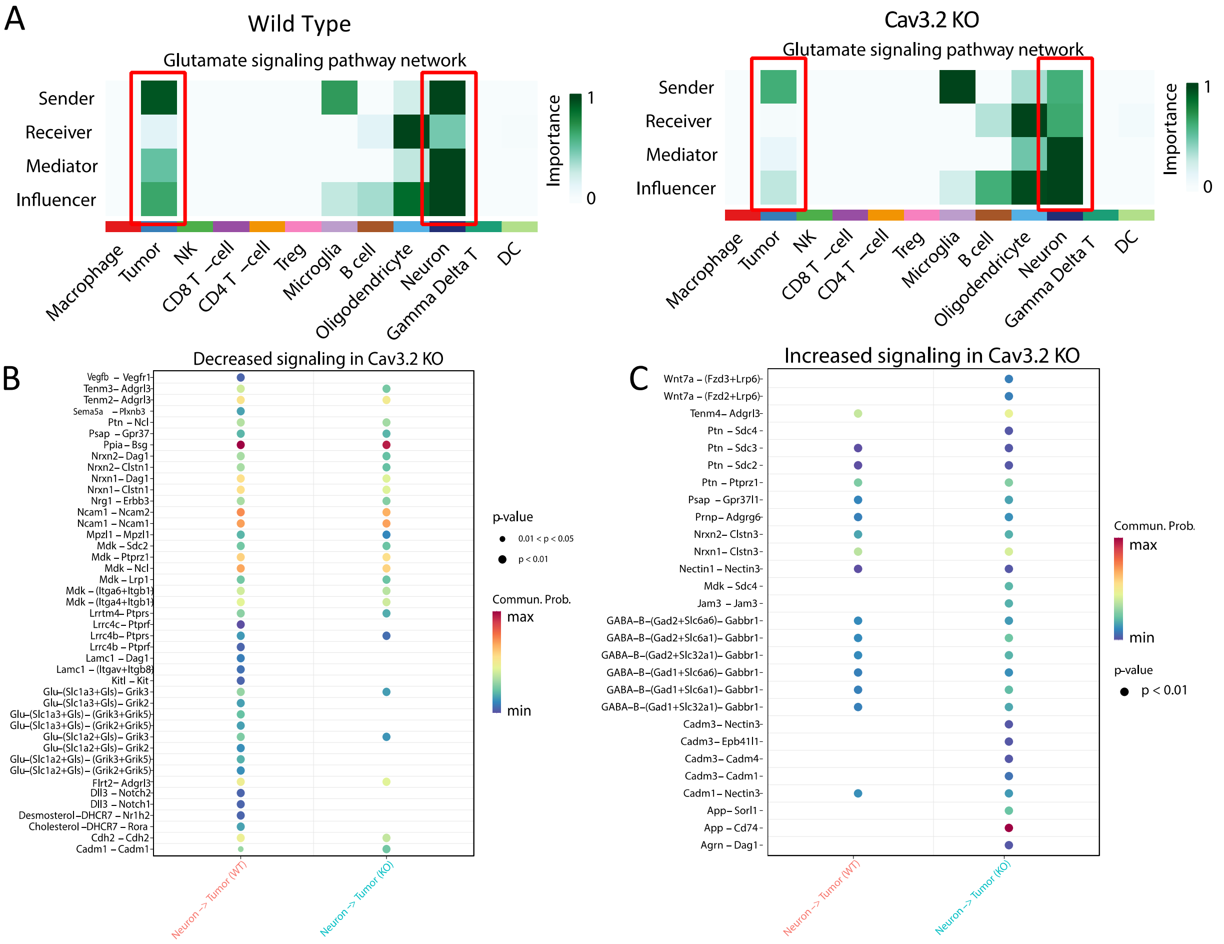


**Supplemental Figure 6:** Microenvironmental Cav3.2 regulates glutamate release. A) Heatmap of Glutamate signaling pathway network in WT and Cav3.2 KO tumors showing a decrease in glutamate signaling in the Cav3.2 KO tumors and neurons. B) Analysis of decreased signaling pathways from neurons to tumors in WT and Cav3.2 KO. The red boxes highlight glutamate and neuron related signaling pairs. C) Dotplot of increased signaling in Cav3.2 KO tumors from neurons to tumor cells.

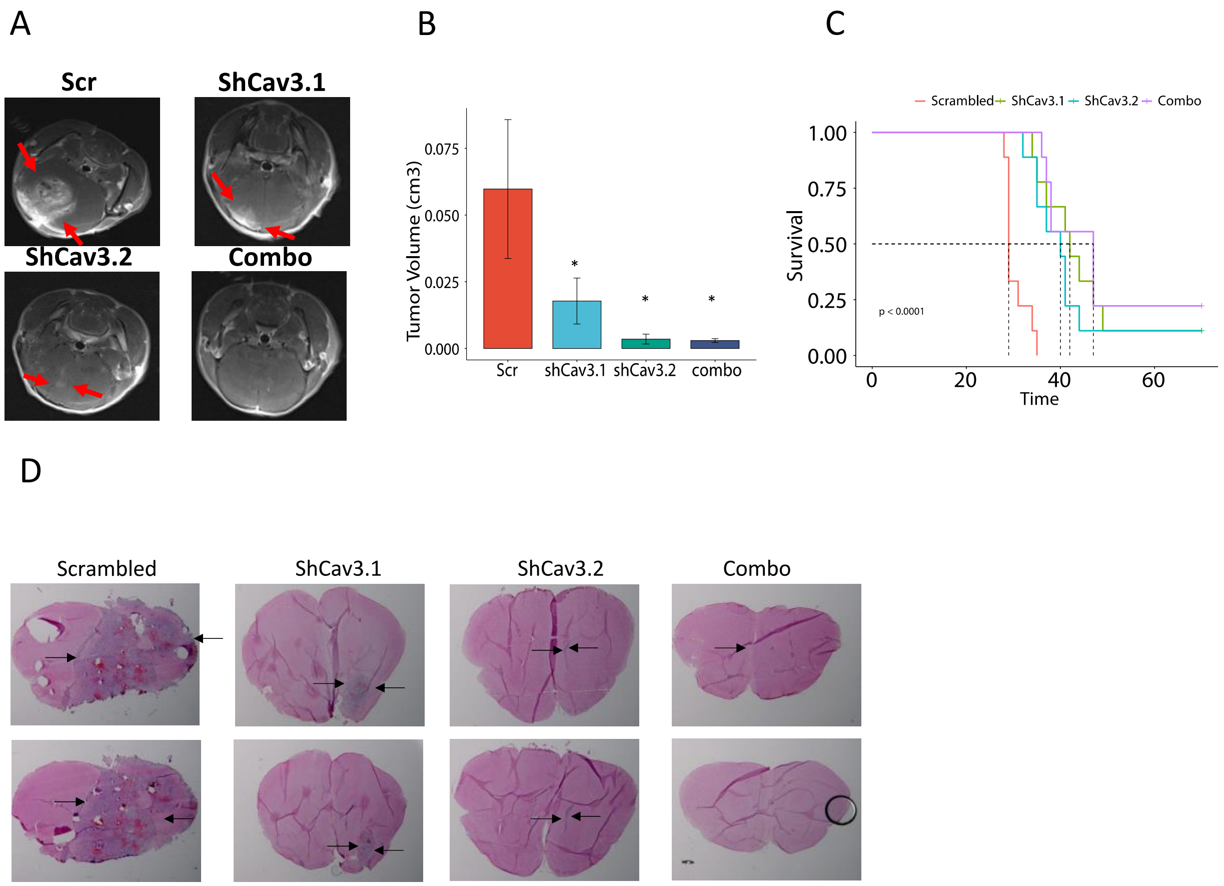


**Supplemental Figure 7:** A) Representative MRI of RCAS/TVA mouse models with Scr, ShCav3.1, Sh-Cav3.2, Combo [50]. B) Quantification of tumor volume in the RCAS/TVA mouse model. C) Kaplan Meier survival curves of the RCAS/TVA mouse model. D) H&E staining of RCAS/TVA mouse model for Control group, shCav3.1, shCav3.2 and combo (shCav3.1+ shCav3.2).

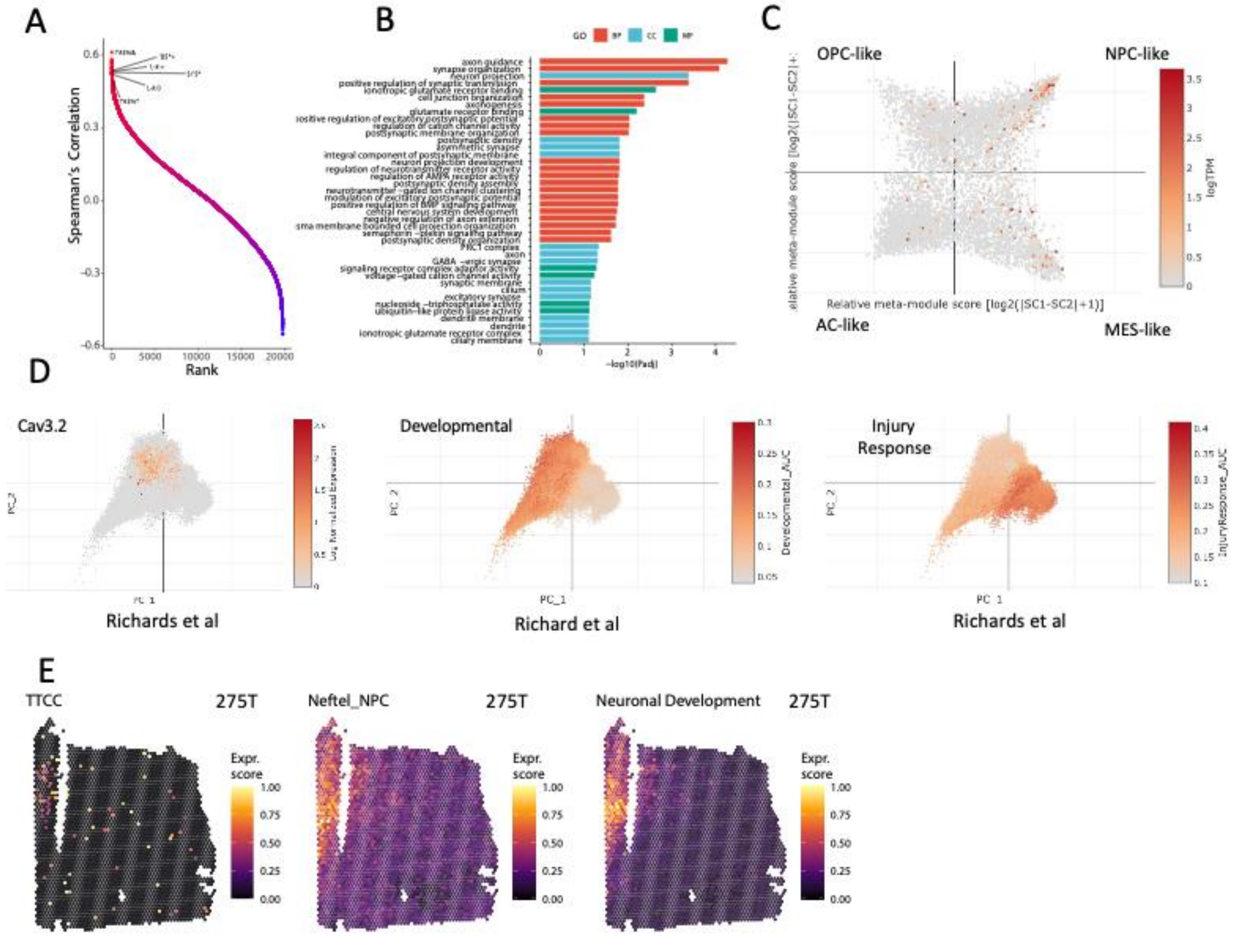


**Supplemental Figure 8:** T-type calcium channels are expressed with neuronal genes. A) Spearmen correlation top genes correlated with Cav3.2 within GBM TCGA dataset. B) Gene ontology of Biological pathways (BP), Cellular Compartment (CC), Molecular Function (MF) of the top positive correlated genes with Cav3.2 in GBM TCGA dataset. C) Expression of Cav3.2 in Neftel dataset showing enrichment within NPC-like cells. D) Expression scores of spatial transcriptomics 275T tumor of T-type calcium channels (Left), Neftel NPC (Middle), Neuronal Developmental Signature (Right). E) Expression of Cav3.2 (Left) in the Richards dataset which overlaps with Developmental cells (middle) compared to injury response (right).
